# Can calmodulin bind to lipids of the cytosolic leaflet of plasma membranes?

**DOI:** 10.1101/2023.12.12.571079

**Authors:** Federica Scollo, Carmelo Tempra, Hüseyin Evci, Miguel Riopedre-Fernandez, Agnieszka Olżyńska, Matti Javanainen, Arunima Uday, Marek Cebecauer, Lukasz Cwiklik, Hector Martinez-Seara, Pavel Jungwirth, Piotr Jurkiewicz, Martin Hof

**Affiliations:** J. Heyrovsky Institute of Physical Chemistry of the Czech Academy of Sciences, Dolejškova 2155/3, Prague 8, 182 23, Czech Republic; Institute of Organic Chemistry and Biochemistry of the Czech Academy of Sciences, Flemingovo nam. 2, Prague 6, 166 10, Czech Republic; Department of Chemistry, Faculty of Science, University of South Bohemia in Ceske Budejovice, 370 05 Ceske Budejovice, Czech Republic; Institute of Biotechnology, University of Helsinki, 00790 Helsinki, Finland

## Abstract

Calmodulin (CaM) is a ubiquitous calcium-sensitive messenger in eukaryotic cells. It was previously shown that CaM possesses an affinity for diverse lipid moieties, including those found on CaM-binding proteins. These facts together with our observation that CaM accumulates in membrane-rich protrusions of HeLa cells upon increased cytosolic calcium, motivated us to perform a systematic search for unmediated CaM interactions with model lipid membranes mimicking the cytosolic leaflet of plasma membranes. A range of experimental techniques and Molecular Dynamics simulations proves unambiguously that CaM interacts with lipid bilayers in the presence of calcium ions. Lipids phosphatidylserine (PS) and phosphatidylethanolamine (PE) hold the key to CaM-membrane interactions. Calcium induces an essential conformational rearrangement of CaM, but its binding to the headgroup of PS also neutralizes the membrane negative surface charge. More intriguingly, PE plays a dual role - it forms hydrogen bonds with CaM, but also destabilizes the lipid bilayer to increase exposure of hydrophobic acyl chains to the interacting proteins. Our findings suggest that upon increased intracellular calcium concentration, CaM and the cytosolic leaflet of cellular membranes can be functionally connected.

## INTRODUCTION

CaM is a multipotent regulator of diverse vital processes in cells. It converts changes in the intracellular calcium concentration to signaling events, specificity of which is determined by its interaction with a variety of proteins. Up to four calcium ions bind to “EF hand” sites (helix–loop– helix motifs) in the two globular domains ^[1]^ inducing conformational rearrangement of CaM to an open, “relaxed” structure (holo-CaM). This is associated with the exposure of flexible hydrophobic pockets, which are involved in the binding of Holo-CaM to a specific set of signaling proteins ^[2]^. Typically, hydrogen bonds are formed between the two hydrophobic anchors of the amphipathic helices in target proteins and a hydrophobic pocket of holo-CaM. ^[3]^ The large variability of binding motifs, however, indicates extensive flexibility of CaM to accommodate diverse binding partners.

Since several targets of holo-CaM are membrane proteins, ^[3]^ it is tempting to speculate that membranes play a role in CaM regulation or form a platform for the complex activity of this highly promiscuous protein. Earlier observations ^[4-6]^ indicated that increased expression of CaM in liver cells can cause transient decrease in membrane fluidity upon increase in intracellular calcium concentration. Whether CaM directly affects lipids in the membrane or the effect is mediated via CaM-binding proteins has not been addressed. Recent studies demonstrated that holo-CaM binds lipid moieties on several CaM-binding proteins. ^[7-9]^ For example, rising intracellular calcium concentration induces sequestration of the prenyl lipid moiety (farnesyl) of a small, highly oncogenic GTPase – KRas4b to the hydrophobic pocket of the C-terminal lobe in holo-CaM. ^[10]^ Moreover, Kovacs and co-workers demonstrated a direct interaction between CaM and a lipid, sphingosylophosphorylcholine (SPC), in the absence of other proteins. ^[8]^ SPC competed with CaM-target proteins indicating a regulatory role of this secondary messenger in CaM function. ^[8]^ Similar effect was observed for related lipids: sphingosine, galactosylsphingosine and glucosylsphingosine. ^[11]^ However, these results were obtained either for lipid monomers or micelles and do not address the interaction of CaM with membranes. Nevertheless, they suggest that CaM owns the capacity to directly bind lipids, which opens the possibility that CaM actively uses cellular membranes for its function.

In this work, we address the question whether CaM can bind to model lipid membranes mimicking the cytosolic leaflet of plasma membranes. Specifically, we studied the calcium dependent interaction of CaM with lipid bilayers using a panel of membrane models and methods, i.e. confocal fluorescence microscopy, fluorescence correlation spectroscopy (FCS), surface plasmon resonance (SPR), generalized polarization (GP) and molecular dynamics (MD) simulations. All data demonstrate the ability of holo-CaM to associate with membranes containing both phosphatidylethanolamine and phosphatidylserine in the absence of other proteins.

## RESULTS AND DISCUSSION

In live-cell imaging experiments, we observed increased presence of CaM tagged with EGFP (EGFP-CaM) in membrane-rich protrusions of HeLa cells treated with ionomycin (**Figure 1A** and Supplementary **Figure S1**), which rapidly increases intracellular calcium concentration. In untreated cells, little to no signal of EGFP-CaM was detected in membrane protrusions. These results suggest that CaM approaches membranes upon Ca^2+^ intake. But, is this solely due to CaM interaction with membrane proteins or it can directly associate with lipid bilayers? What would be the consequences of such association? To untangle this possibility and the role of specific lipids in a putative CaM-membrane interaction, we prepared several model membrane systems with the following compositions: PC, PC/PS (8:2), PE/PS/CH (6:2:2), PE/PC/CH (6:2:2), PE/PS/PC (6:2:2), PC/PS/CH (6:2:2) and PE/PC/PS/CH (4:2:2:2) reported in **Table 1**, where PC, PE, PS and CH stand for 1-palmitoyl-2-oleoyl-sn-glycero-3-phosphatidylcholine, 1,2-dioleoyl-phosphatidylethanolamine and1,2-dioleoyl-phosphatidylserine and cholesterol, respectively (see **Figure S1** for the chemical structures).

**Table 1.** Compositions of model lipid membranes used in this study.

| Abbreviation | POPC (mol%) | POPS (mol%) | POPE (mol%) | Cholesterol |
| --- | --- | --- | --- | --- |
| PC [a, b] | 100 | - | - | - |
| PE [b] | - | - | 100 | - |
| PC/PS [a, b] | 80 | 20 | - | - |
| PE/PS [b] | - | 20 | 80 | - |
| PE/PC [b] | 20 | - | 80 | - |
| PC/CH [b] | 80 | - | - | 20 |
| PE/CH [b] | - | - | 80 | 20 |
| PC/PS/CH [a, b] | 60 | 20 | - | 20 |
| PE/PS/CH [a, b] | - | 20 | 60 | 20 |
| PE/PC/CH [a, b] | 20 | - | 60 | 20 |
| PE/PS/PC [a] | 20 | 20 | 60 | - |
| PE/PC/PS/CH [a] | 20 | 20 | 40 | 20 |

[a] Used in the experiments [b] used in the simulations.

**Figure 1.**
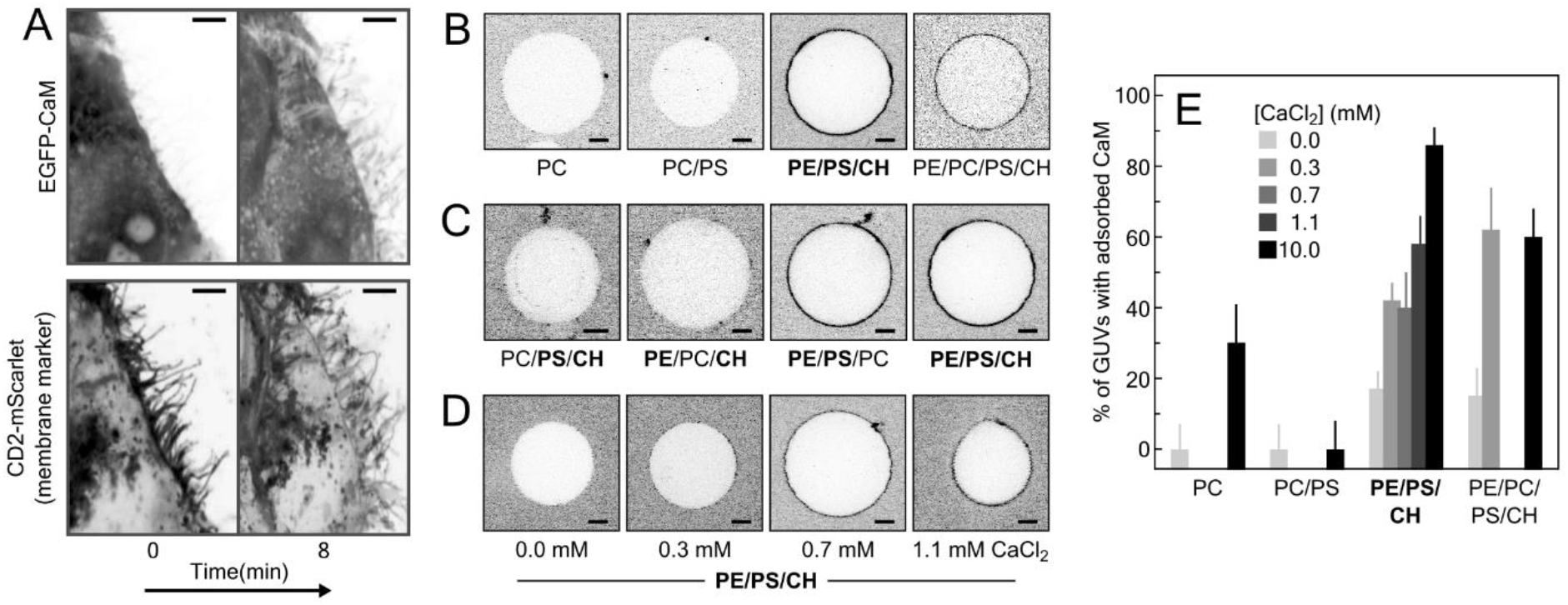
a) Fluorescence confocal images of transfected HeLa cells expressing EGFP-CaM (upper images) a membrane marker CD2-mScarlet (lower images). Relocalization of EGFP-CaM to membrane protrusions is visible after ionomycin treatment. Scale bars of 5 μm are shown in the right upper corners. b) and c) Representative cross sections of GUVs of different lipid compositions (see Table1 for details) incubated with 30 nM CaM-R solution at 10 mM CaCl_2_. d) Representative cross-sections of PE/PS/CH and 30 nM CaM-R at different CaCl_2_ concentrations: 0, 0.3, 0.7 and 1.1 mM. All the images were corrected for background intensity. 5 μm scale bars are shown in the lower right corners. N > 9 (≥ 3 different electroformations). e) Percentage of GUVs showing CaM-R adsorption on their membranes for PC, PC/PS, PE/PS/CH and PE/PC/PS/CH at different CaCl_2_ concentrations.

First, we studied the adsorption of rhodamine-B labeled CaM (CaM-R) to giant unilamellar vesicles (GUVs) using confocal fluorescence microscopy.

**Figure 1B** demonstrates CaM-R adsorption to PE/PS/CH and PE/PC/PS/CH, but not to PC nor PC/PS vesicles (GUVs). Since the adsorption was the most pronounced at PE/PS/CH membranes, we investigated which of its components promotes the effect by replacing one at a time with PC.

From the resulting lipid compositions: PC/PS/CH, PE/PC/CH, and PE/PS/PC only the last one was effective (**Figure 1C**). It is thus evident that both PE and PS were required for the CaM-R adsorption, while cholesterol was found to be dispensable. All these experiments were performed in the presence of 10 mM CaCl_2_ however, much lower concentrations were sufficient for the adsorption (**Figure 1D, E**). CaM-R adsorption to PE/PS/PC was detectable already at 0.3 mM CaCl_2_.

The observed CaM adsorption required rather long incubation times of > 4 h, which contrasts with live-cell data. To approach earlier stages of the CaM-lipid membrane interactions, we took advantage of the sensitivity of fluorescence correlation spectroscopy (FCS). Fluorescence fluctuations caused by CaM diffusion were measured in the bulk (solute) and at the surface of GUVs (see Methods in the SI for more details). The FCS data indicate that even in the absence of PE and Ca^2+^ there was a minute fraction of CaM at the surface of the GUV membrane in < 30 min (**Figure 2A**). However, the presence of PE, PS and CaCl_2_ increased the fraction of the CaM associated with membranes (**Figure 2A**).

**Figure 2.**
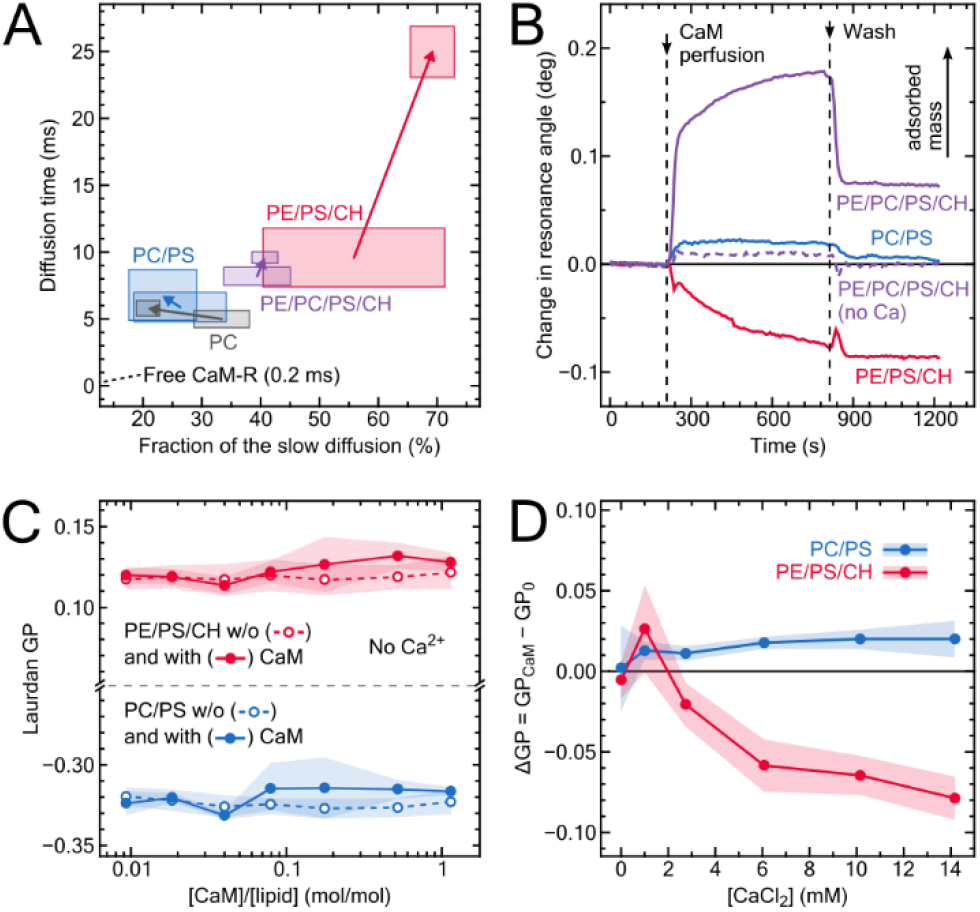
a) Longer FCS diffusion time as a function of its fraction measured for 30 nM CaM-R diffusing at the surface of giant vesicles (GUVs) in the absence and in the presence of 10 mM CaCl_2_ (increasing CaCl_2_ concentration depicted by the arrows). Rectangles represent SD, N > 10. b) Change in the resonance angle measure using surface plasmon resonance for supported lipid bilayers (SPB) upon perfusion with 0.6 μM CaM solution. The solution contained 10 mM CaCl_2_ (solid lines) or no calcium (dashed line). Lipid composition of SPBs given in **Table 1**. Curves represent individual measurements. c) Generalized polarization (GP) of Laurdan in extruded liposomes (LUVs) as a function of CaM/lipid ratio in the absence of calcium. Open symbols and dashed lines represent control experiments without CaM. d) ΔGP induced by CaM as a function of CaCl_2_ concentration for CaM/lipid ratio of 1.1. ΔGP is defined as GP for CaM-containing sample – GP of a CaM-free sample. All measurements were performed at 37°C. Error bands represent SE (N=3). Lipid compositions of all the model systems shown in this figure are given in **Table 1**.

To avoid the impact of the fluorophore, we next used surface plasmon resonance (SPR) – a label-free technique ^[12-15]^ enabling determination of real-time protein adsorption to the surface of lipid membranes. ^[16-17]^ Supported lipid bilayers (SPBs) were created by vesicle deposition ^[18]^ on a gold two-channel SPR sensor covered with SiO_2_. Perfusion with 0.6 μM unlabeled CaM with 10 mM CaCl_2_ resulted in significant adsorption of CaM on PE/PC/PS/CH, which was stable even after subsequent washing with CaM-free buffer (**Figure 2B**). The adsorption was detectable within a minute and increased during the period of the measurement (10 min). In the absence of calcium or PE, no CaM adsorption was detected. Increased PE content in this experiment resulted in membrane disintegration, as shown by the surface mass loss for PE/PS/CH in **Figure 2B**. In analogy, while performing confocal microscopy, we observed GUV rupture after CaM-R addition, to PE/PS/CH GUVs (≥40% of vesicles). This indicates that CaM-R can strongly perturb PE-containing membranes, up to the point where their integrity becomes compromised. Calcium-dependent binding of negatively charged proteins to PS-containing membranes is a well-known phenomenon. ^[19-20]^ However, how can one rationalize that PE is essential for holo-CaM binding to lipid membrane? Calcium ions induce conversion of apo-CaM into holo-CaM exposing its hydrophobic pockets. As described before, ^[7-10]^ a hydrocarbon tail of phospholipids can bind into these pockets. In membranes, however, the lipid tails are normally kept hidden in the bilayer core and not exposed to potential binding partners. Thus, CaM would need to surpass lipid-lipid interactions to access lipid acyl chains. We suggest that this process can be facilitated by PE – a cone-shaped lipid prone to destabilize lamellar structure of a lipid membrane. ^[21]^

Confocal microscopy and SPR experiments indicated that holo-CaM can disintegrate PE/PS/CH membranes, which implies changes in the structural and dynamical properties of the lipid bilayer. To monitor such changes in membrane properties in the presence of CaM and calcium, we used generalized polarization (GP) of Laurdan. ^[22]^ This method probes the bilayer fluidity, more specifically the lipid mobility at the glycerol backbone level. ^[23]^ It is exceptionally sensitive to the alterations of membrane physical properties,^[24]^ including those caused by proteins and peptides. ^[25-27]^ In the absence of calcium, no effects of CaM on PC/PS and PE/PS/CH large unilamellar vesicles (LUVs) were detected (**Figure 2C**). In the presence of increasing calcium concentration, CaM caused a significant decrease in Δ*GP* of PE/PS/CH LUVs at 3 mM CaCl_2_ and above. Noteworthy, a decrease in membrane rigidity represented by a decrease of the GP value upon the binding of any other protein has not been reported in earlier studies. We suggest that membrane destabilization caused by CaM-PE interaction is the reason for the lowering of Laurdan Δ*GP* observed here for PE/PS/CH in the presence of Ca^2+^ (**Figure 2D**). In the case of PC/PS membranes, a small, but statistically significant elevation of Δ*GP*, can be observed already at 1 mM CaCl_2_, and it does not further increase at larger Ca^2+^ concentrations. In analogy to our previous work, ^[25, 27-30]^ this mild rigidification of the lipid bilayer might suggest a peripheral association of CaM with the lipid bilayer. For CaM, we discovered the strongest calcium-dependent alteration of the headgroup mobility for the PE containing bilayers, but in a unique direction (**Figure 2D**). Evidently, CaM perturbs the cohesive interactions between the lipid molecules, which is consistent with the disruptive interactions observed between CaM and PE/PS/CH membranes.

The different experimental approaches described above reveal three components that favor the adsorption of CaM to lipid bilayers: PE and PS lipids as well as calcium ions. To better understand the molecular nature of the initial step in CaM adsorption and to get insights into the role of these three components, we ran atomistic multi-microsecond MD simulations of CaM at membranes with the lipid compositions used in the experiments, and the following additional ones: PE, PE/PS (8:2), PE/PC (8:2), PC/CH (8:2), and PE/CH (8:2) (see **Table 1**, Experimental Procedure and **Table S2** in the SI for a detailed description of the systems). Briefly, CaM was initially placed in the aqueous phase, distant from the studied membranes, each of which had different compositions and each containing a total of 200 lipids. Then, CaM was allowed to spontaneously interact with the membrane during a 5 μs-long simulation. To model the interactions among charged species, we used the implicitly polarizable simulations models that have captured the calcium-bridged bridging of PKC*α*-C2 domain onto PS lipids in our earlier work (see SI for details). ^[31]^ Contact probability of membranes with apo- and holo-CaM (4 calcium ions bound to the calcium-binding sites) at different CaCl_2_ concentrations was calculated from the simulations. The data are represented in **Figure 3A**.

**Figure 3.**
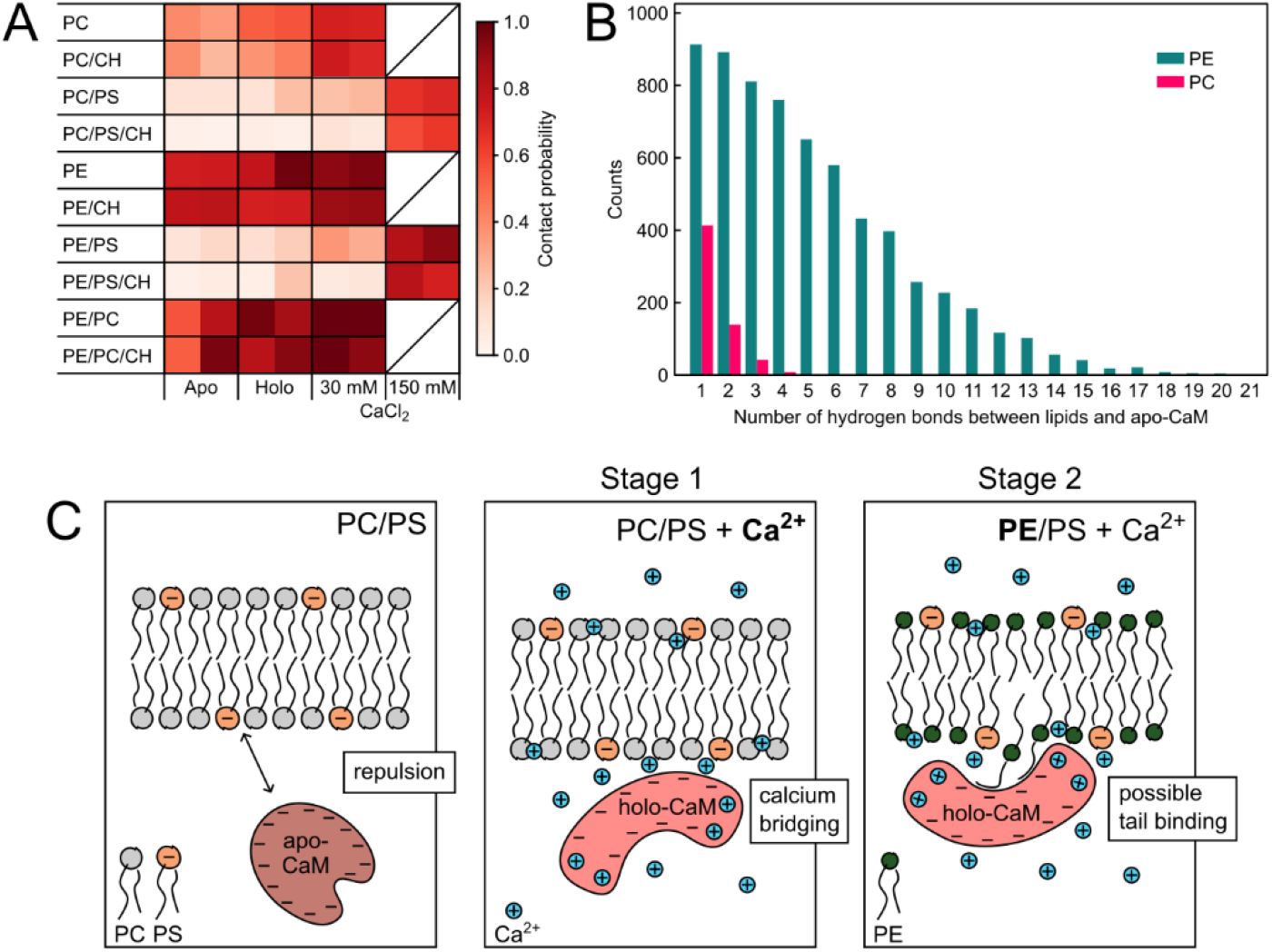
a) Probability of contact between CaM and lipid membranes. Rows represent different lipid compositions (see **Table 1** for details). Columns represent different CaM forms and CaCl_2_ concentrations. Each cell contains two different replicates of the simulations (each 5 μs long). Details of the simulations are provided in **Table S1** in the SI. b). Histograms of hydrogen bonds between apo-CaM and membranes for PC and PE membranes. c) Molecular sketch illustrating our hypothesis of two-stage model of CaM binding to lipid bilayer. In the absence of calcium ions apo-CaM is electrostatically repelled from the negatively charged surface of PC/PS membrane (left sketch). Calcium causes formation of holo-CaM exposing its hydrophobic binding pocket. Calcium ions also adsorb to negatively charged surfaces of holo-CaM and PC/PS membrane bridging them (stage 1, middle sketch). This process can cause transient association of CaM with the membrane. PE forms hydrogen bonds with holo-CaM, which further attracts the protein to the membrane. PE also destabilizes the membrane, increasing the probability of exposure of hydrophobic lipid tails and their binding to the hydrophobic pocket of CaM (stage 2, right sketch). This could lead to membrane rearrangement and stable CaM adsorption or to destruction of the membrane.

In agreement with the experiments, MD simulations demonstrate that exchanging PE with PC in tested membranes lowered the probability of their contact with CaM (apo-CaM and holo-CaM at any calcium concentration). This difference can be explained at the molecular level by a higher capability of PE to form hydrogen bonds with the protein (**Figure 3B**). Interestingly, the contact probabilities in the absence of calcium ions in solution drop dramatically once PS is added to the lipid mixtures, likely because of the net negative charge of PS, which repels CaM, a highly negatively charged protein, and prevents its contacts with the membrane. However, when Ca^2+^ concentration is high, contact probability of CaM with PS-containing membranes significantly increases, i.e., to about 80% at 150 mM of calcium (**Figure 3A**). Indeed, the contact probability of CaM with membranes increases with calcium concentration to different extent but consistently for all membrane compositions, indicating that calcium ions are key for the protein to approach the membrane (**Figure 3A**). While this concentration of calcium seems high compared to the experiments, the simulations actually contain a modest total number of ions, and in experiments its local concentration at the vicinity of charged membranes might be similarly elevated. We observed similar calcium-dependence for CaM presence in membrane-rich protrusions, but cannot exclude protein mediators in live-cell experiments (**Figure 1A**). The presence of cholesterol appears dispensable for the protein-membrane binding for all the simulated compositions, confirming the experimental results **Figures 1-2**. Additional simulations show that for systems containing PE and Ca^2+^, binding modes with more EF loops bound at the same time occur (**Figure S4**). The data suggest that the probability that CaM lies down on the membrane surface are increased by the presence of PE.

Based on our results and on previous studies, ^[8-9]^ we propose a two-stage model of CaM-membrane interactions, illustrated in **Figure 3C**. As demonstrated before, adsorption of Ca^2+^ to PC/PS membrane can neutralize its negative charge, or even turn it positive (in case of overbinding ^[32]^). Our MD simulations indicate that Ca^2+^ bridging might be the driving force for initial CaM interactions with PS-containing membranes (Stage 1, **Figure 3C**, middle panel). This process can lead to a transient association of CaM with the membrane, which we consider as the first necessary stage for the binding to occur. PE likely plays a dual role: 1) its ability to form hydrogen bonds with the holo-CaM can accelerate CaM attraction to the membranes, but 2) PE, with its preference for non-lamellar structures destabilizes lipid bilayers, which can lead to the exposure of lipid tails to the hydrophobic pocket of holo-CaM (Stage 2, **Figure 3C**). Such events may lead to a more stable interaction of CaM with membranes.

What makes the interaction of CaM with membranes mimicking the cytosolic leaflet of cellular membranes unique? Ca-dependence of protein association with membranes is a well described phenomenon (e.g., for C2 domain of phospholipase A2 ^[33]^). Anionic phospholipids regulate function of numerous membrane-associated signaling molecules (e.g., Akt ^[34]^). However, none of these Ca and/or lipid-dependent proteins exhibits such an interplay between PE and PS.

In general, a direct regulation of proteins by PE is less well understood than by PS. PE has indirect positive effect on the interactions of proteins with negatively-charged membranes. ^[35-36]^ This is especially pronounced at lower content of PS. In the case of coagulation factors, PS and PE functioned in synergy: PS was sufficient for a protein binding to membranes, but the presence of PE reduced K_D_ values of this interaction.^[35-36]^ Similar positive effect of PE was observed for protein binding to DAG-containing membranes (e.g., protein kinase C ^[35]^). PE and PS are metabolically closely related, which can be also reflected in their functions. ^[37]^ We believe that in the case of holo-CaM binding to membranes, PE and PS might as well work synergistically. Lanthionine-containing peptide antibiotics, duramycin and cinnamycin, bind to lipid bilayers in a strictly PE-dependent manner. ^[38]^ At high PE concentration, these peptides destabilize membranes and cause cell death. ^[39]^ This is indeed similar to our observation that holo-CaM can destabilize PE-containing membranes at higher calcium concentrations. Since the formation of membrane pores, as shown for cinnamycin, is highly improbable, ^[39]^ we hypothesize that holo-CaM may bind to the phospholipids released by the PE-containing membranes or to lipid acyl chains exposed at the surface.

What supports our view about CaM-acyl chain interactions? Prenyl moiety of KRas4b binds to the hydrophobic groove of holo-CaM regulating its function ^[40]^ Similar processes were described for holo-CaM interacting with prenylated RalA ^[7]^ Moreover, holo-CaM binds CAP23/NAP22 protein by accommodating its myristoyl moiety in its hydrophobic pocket.^[41]^ This indicates that alkyls can facilitate holo-CaM binding as efficiently as prenyls. Basic residues in the proximity of the prenylation (myristoylation site) facilitate holo-CaM attraction to the target lipidated proteins and stabilize bimolecular complex. Such scenario resembles our observation that, in the presence of Ca^2+^, PS attracts CaM to the membrane, but PE is required for membrane destabilization and acyl chain exposure for a more stable binding of holo-CaM (**Figure 3C**).

## CONCLUSION

Our study demonstrates that, under certain conditions, CaM, which is normally a hydrophilic, highly negatively charged and well-soluble protein, can interact with model lipid membranes. It revealed specific involvement of PS and PE lipids, which are major components of the cytosolic leaflet of the plasma membrane. Based on our results and the existing literature, we propose a two-stage model of CaM interaction with membranes: 1) Ca^2+^ adsorption to PS headgroups facilitates transient interaction of holo-CaM with a bilayer; 2) PE, by reducing cohesive lipid assembly and the potentially increased probability of acyl chain exposure, further promotes and stabilizes CaM adsorption to membranes (**Figure 3C**). The later can lead to membrane destabilization or even disintegration. While the role of PS follows a typical pattern found for calcium-mediated protein/membrane interactions, PE acts in a unique fashion. Our two-stage binding model of holo-CaM to PS/PE-containing membranes is in line with the Ca-independent adsorption of apo-CaM to SPC micelles, ^[8]^ and expanding it to lipid bilayers. Overall, our study provides a mechanistic insight into the intricate interplay between CaM and membranes, suggesting a possible regulatory role of membrane lipids in the complex activity of CaM.

## ACKNOWLEDGEMENTS

We would like to thank Jan Sýkora for sharing his expertise in several discussions and for arranging SPR measurements, Darina Majovská, Harsha Mavila and Dalibor Pánek (IMCF, BIOCEV, Vestec) for technical assistance with cell culture, DNA cloning and live-cell microscopy, respectively. The authors also gratefully thank Matěj Pastucha, Radka Obořilová, and Zdeněk Farka (Department of Biochemistry, Faculty of Science, Masaryk University, and CEITEC MU, Masaryk University, Brno) for their theoretical and technical expertise in SPR technique, and acknowledge CF Nanobiotechnology of CIISB, Instruct-CZ Centre, supported by MEYS CR (LM2018127). The measurements at the Imaging Methods Core Facility in BIOCEV, Vestec, Czech Republic were supported by MEYS CR grant Z.02.1.01/0.0/0.0/16_013/0001775. We thank CSC– IT Center for Science for computational resources. We are grateful for the financial support provided for this work by the Czech Science Foundation via grant 19-26854X.

## AUTHOR CONTRIBUTION

F.S., H.E., M.R.F., A.O., A.U., P.Jur. carried out the experiments. F.S., H.E., P.Jur. analyzed the experimental data. F.S., C.T., P.Jur., prepared the figures. F.S. and P.Jur. conceived and planned the experiments. F.S., H.E., M.R.F., A.O., A.U., M.C., P.Jur., M.H. contributed to the interpretation of the results. M.C., P.Jur. and M.H. supervised the experiments. C.T. and M.J. carried out and analyzed the simulations. C.T., M.J., L.C., H.M.S. discussed and co-designed the simulations. C.T., M.J., L.C., H.M.S. and P.Jun. contributed to the interpretation of the MD simulations result. H.M.S. and P.Jun. supervised the computational part. F.S., C.T., M.C., P.Jur and M.H. wrote the manuscript. H.M.S. and P.Jur. conceived the original idea. H.M.S., P.Jun., P.Jur. and M.H. supervised the project. All authors contributed to the discussion and provided critical feedback.

