## Supplementary Material for "Can calmodulin bind to lipids of the cytosolic leaflet of plasma membranes?"

**ABSTRACT**

Calmodulin (CaM) is a ubiquitous calcium-sensitive messenger in eukaryotic cells. It was previously shown that CaM possesses an affinity for diverse lipid moieties, including those found on CaM-binding proteins. These facts together with our observation that CaM accumulates in membrane-rich protrusions of HeLa cells upon increased cytosolic calcium, motivated us to perform a systematic search for unmediated CaM interactions with model lipid membranes mimicking the cytosolic leaflet of plasma membranes. A range of experimental techniques and Molecular Dynamics simulations proves unambiguously that CaM interacts with lipid bilayers in the presence of calcium ions. Lipids phosphatidylserine (PS) and phosphatidylethanolamine (PE) hold the key to CaM-membrane interactions. Calcium induces an essential conformational rearrangement of CaM, but its binding to the headgroup of PS also neutralizes the membrane negative surface charge. More intriguingly, PE plays a dual role - it forms hydrogen bonds with CaM, but also destabilizes the lipid bilayer to increase exposure of hydrophobic acyl chains to the interacting proteins. Our findings suggest that upon increased intracellular calcium concentration, CaM and the cytosolic leaflet of cellular membranes can be functionally connected.

### Experimental Procedures

#### Materials

Bovine brain calmodulin of >95% purity was obtained from Sigma-Aldrich (St.Louis, MO). Porcine brain calmodulin >98% purity was obtained from Analytik Jena Roboscreen GmbH (Leipzig, Germany). Wheat (Triticum aestivum) calmodulin fluorescently labeled with rhodamine B (CaM-R) of 98% purity was purchased from rPeptide (Watkinsville, GA, USA). Lipids: 1-palmitoyl-2-oleoyl-*sn*-glycero-3-phosphocholine (POPC), 1-palmitoyl-2-oleoyl-*sn*-glycero-3-phospho-L-serine (POPS), 1-palmitoyl-2-oleoyl-*sn*-glycero-3-phosphoethanolamine (POPE), 1-palmytoil-2-oleoyl-*sn*-glycero-3-phosphoethanolamine-N-(cap biotinyl) (DOPE-cap-biotin); and cholesterol (ovine wool, 98%) were purchased from Avanti Polar Lipid (Alabaster, AL, USA). Fluorescent dyes: 6-dodecanoyl-N,N-dimethyl-2-naphthylamine (Laurdan) and rhodamine B were purchased from Molecular Probes (Eugene, OR, USA) and Sigma-Aldrich, respectively. Potassium hydroxide, potassium, calcium chlorides, 4-(2-hydroxyethyl)-piperazine-1-ethanesulfonic acid sodium salt (HEPES), ethylene glycol-bis(β-aminoethyl ether)-N,N,N′,N′-tetraacetic acid (EGTA), Bovine Serum Albumin (BSA), sucrose, D-(+)-glucose, Dulbecco`s Modified Eagle Medium (DMEM), and Fetal Bovine Serum (FBS) were purchased from Sigma-Aldrich (St.Louis, MO; all chemicals 99% purity, unless otherwise stated). Aqueous solutions were prepared in calcium free water (Sigma-Aldrich, St.Louis, MO), nuclease free water or milliQ water. Spectroscopic grade chloroform, methanol and ethanol (purity 96%) were purchased from Merck (Darmstadt, Germany). Biotinylated bovine serum albumin (biotin-BSA) and Indo-1 pentapotassium salt, were purchased from Thermo Fisher (Waltham, MA USA). Streptavidin was purchased from IBA Lifesciences (Göttingen, Germany). Restriction enzymes kit, Antarctic Phosphatase treatment kit, Polymerase Chain Reaction (PCR) kit, and ligation kit were purchased from New England Biolabs (Ipswich, MA, USA). QIAGEN Miniprep kit and QIAGEN Midiprep kit were purchased from QIAGEN (Hilden, Germany). M-Slide 8 Well ibiTreat and glass bottom #1.5 coverslip were purchased from ibidi (Graefelfing, Germany). TransIT-X2® Dynamic Delivery System were purchased from Mirus Bio LLC (Madison, WI, USA).

#### DNA cloning

The generation of EGFP-calmodulin (CaM-EGFP) fusion protein construct involved a series of processes. The calmodulin construct was obtained in plasmid pEX-A128 (Eurofins Scientific, Nantes, France). Calmodulin was extracted and introduced to the vector plasmid pXJ41 by Restriction Enzyme Digestion at the BamHI and XhoI restriction sites. The digested vector pXJ41 and the inserted calmodulin were subjected to ligation and then transformation into Top10 E. coli bacteria cells. Selected transformed colonies were subjected to Colony Polymerase Chain Reaction (PCR) to identify successfully transformed colonies. The plasmids were then isolated from these colonies using a Qiagen Miniprep Kit (QIAGEN, Hilden, Germany). These plasmids were sequenced and stored. The EGFP construct was amplified by PCR from a pXJ41 plasmid containing CD2-EGFP using GFP-EcoRI forward and GFP-BamHI reverse primers (Sigma-Aldrich, St.Louis, MO, USA). This EGFP construct was introduced into the sequenced pXJ41 plasmid with calmodulin by Restriction Enzyme Digestion at EcoRI and BamHI restriction sites. This was followed once again by ligation, transformation and Colony PCR. The plasmids were extracted using Qiagen MidiPrep Kit before sequencing and stored for further experiments.

**HeLa cell culture**

HeLa cells (ATCC) regularly tested for surface markers, morphology and contamination were grown in DMEM (High-Glucose, Sigma-Aldrich) supplemented with 10% FBS (Gibco) in the Eppendorf® Cell Culture Flask T-25 (TC treated, with filter cap) in a humidified incubator at 37 °C and 5% $CO_2$ (Eppendorf). Before transfection, 15 000 cells were seeded in a well of μ-Slide 8 Well ibiTreat #1.5 chamber. Next day, cells were transfected with total 1 μg DNA plasmid (0.5 μg DNA of pXJ41-EGFP-CaM and 0.5 μg DNA of pXJ41-CD2-mScarlet) per well using TransIT-X2® Dynamic Delivery System (Mirus) according to the manufacturer's recommendations. Cells were incubated for another 16-24 hours before imaging.

**Live cell imaging and data processing**

Imaging of live cells expressing CaM-EGFP and CD2-mScarlet was performed on a Nikon CSU-W1 equipped with stage-top heating/cooling incubator for live cell imaging. It is based on an inverted widefield microscope Nikon Eclipse Ti2 equipped with motorized XY stage, Perfect Focus System, semi-motorized DIC, LED transmitted light source, multicolor LED for wide-field epifluorescence 60× objective (CF Plan Apo VC 60XC WI, NA = 1.2 water immersion). For excitation, epifluorescence LED light source (488 nm and 561 nm) were used. Fluorescence was detected in a PRIME BSI (Teledyne Photometrics) camera with selective dichroic mirrors and blocking filters. NIS-Elements software was used to operate the microscope and for image acquisition.
Imaging of live cells expressing CaM-EGFP and CD2-mScarlet for quantitative analysis was carried out on a modified Olympus Fluoview 1000 setup (Olympus) equipped with the environmental chamber (OKO-Lab) to keep cells at 37°C and 60× objective (UplanSApo, NA = 1.2 water immersion; Olympus). For excitation, 488 nm solid state laser (Sapphire, 20mW; Coherent) and 532 nm high power pulsed diode laser (TopTica, 10mW; PicoTA) were used. Fluorescence was detected in a single channel mode on PMT detectors equipped with selective dichroic mirrors and blocking filters (Olympus). Transmitted light detector was employed for a brightfield imaging. FluoView 1000 software (Olympus) was used to operate the microscope and for image acquisition. Images were taken before and after addition of ionomycin and waiting for 2 hours to open $Ca^{2+}$ channels. The experiments were performed in less than 24h after transfection to avoid cytotoxic effect of CaM overexpression. [1]
Images were analyzed using ImageJ software. To better visualize the results, minor brightness adjustments were applied. For quantitative analysis of CaM accumulation, the protrusions were first selected in the mScarlet channel (CD2). Afterwards, maximum signal in the EGFP channel (IEGFP(p)) was determined for the selected areas. Similarly, maximal signal in the EGFP channel (IEGFPI) was determined for the cell body (cytosol) of the same cell. CaM accumulation was calculated as a ratio of the maximum signal from protrusions to cytoplasm (IEGFP(p)/IEI(c)) to normalize measured values in cells with varying protein expression levels. At least 7 protrusions per cell were selected. 886 protrusions before and 686 after the treatment with 200 nM ionomycin were analyzed.

**Preparation of model membranes**

Appropriate lipid compositions were prepared by mixing chloroform solutions of lipids and methanol solution of Laurdan (2 mol%, when needed). Organic solvents where then evaporated under stream of nitrogen and vacuum (>60 min). The dry lipid film formed on the electrodes (for electroformation of Giant Unilamellar Vesicles, GUVs) or in the glass vial (for Large Unilamellar Vesicles, LUVs, or Supported Lipid Bilayer, SLBs) was hydrated with an appropriate solution. The desired liposomes were obtained with an appropriate method, each of them described below.

LUVs were formed by extrusion of multilamellar vesicles (MLVs), obtained by 4 min vortexing of dry lipid film hydrated in warm buffer (10 mM HEPES, 150 mM KCl, 0.1 mM EGTA, pH=7.4), through polycarbonate filters (50 cycles, pore diameter 100 nm, Nuclepore, Pleasanton, CA) mounted in a mini-extruder (Avestin, Ottawa, ON, Canada).

Supported lipid bilayers (SLBs) were created directly on an SPB gold sensor covered with $SiO_2$ layer (BioNavis, BN-M0X-2007-001) from Small Unilamellar Vesicles (SUVs) dispersion, in turn obtained by ultrasonicating MLVs dispersion (10 min, 40% power) with subsequent centrifugation (16'000 rpm, 20 min). The sensor was carefully cleaned prior each measurement by using the procedure described elsewhere. [2] SUVs with 1 mM total lipid concentration, buffer (10 mM HEPES, 150 mM KCl, 0.1 mM EGTA, pH=7.4) and 2mM $CaCl_2$ solution were added onto the freshly cleaned sensor and incubated for 1h or more (according to the composition) immediately before SPR measurements.

Giant unilamellar vesicles (GUVs) were prepared by the electroformation method (Angelova et al. 1992) using either titanium plates or platinum wires as electrodes. 100 nmol of lipids (in total) were uniformly spread on two electrodes using Hamiltonian syringe. 1 mol% of DPPE-cap-biotin was present in all GUV formulations for further immobilization of GUVs. Lipid-coated titanium plates were assembled and sealed with parafilm forming a closed chamber that was filled with sucrose solution (osmolarity of 300 $mOsm \cdot kg^{-1}$). Sinusoidal voltage of 10 Hz frequency rising from 0.15 to 1.1 V was applied to the plates for 45 min, then kept at 1.1 V for 2.5 h, and finally raised to 1.3 V at 4 Hz for 30 min. Alternatively, platinum electrodes were placed into Teflon chamber and a sinusoidal voltage of 4 V was applied at 10 Hz for 50 min and then at 2 Hz for 20 min. The electroformation was performed at 45 °C.

### Surface Plasmon Resonance

A MP-SPR Navi 210A (BioNavis, Finland) system consisted of a two-channel experimental design including the detection and reference channels with a parallel injection mode was employed to perform SPR measurements. Degassed running buffer (10 mM HEPES, 150 mM KCl, 0.1 mM EGTA, pH=7.4) was used with a flow rate of 20 μL/min for 10 min, to establish a baseline. After confirming the effective SPB formation and its initial equilibration in the buffer (10 mM HEPES, 150 mM KCl, 0.1 mM EGTA, pH = 7.4), 0.6 μM solution (10 mM HEPES, 150 mM KCl, 0.1 mM EGTA, pH=7.4, 10 mM $CaCl_2$) was injected into the detection channel, whereas the same solution without CaM was injected to the reference channel to rule out possible non-specific binding and compensate for the related nonspecific changes in refractive index. Flow rate of 20 μL/min was used (same as for the running buffer). After the adsorption process was monitored, the sample was flushed with the buffer to check the stability of the adsorbed protein. SPR experiments were performed at 25 °C and using 670 and 785 nm lasers. The surface plasmon resonance angle determined in an angular scan mode was analyzed by using a centroid fitting function, allowing to evaluate the sensor response. The data acquired with two different lasers were averaged. Every curve is presented after normalization and reference subtraction.

**SPBs characterization by Z-Scan Fluorescence Correlations Spectroscopy**

SPBs formed on a gold sensor covered with SiO2 layer (BioNavis, BN-M0X-2007-001) were characterized using fluorescence microscopy and Z-scan fluorescence correlation spectroscopy (Z-scan FCS). All measurements were performed on an inverted confocal microscope, Olympus IX71 (Olympus, Hamburg, Germany), equipped with single-photon counting unit MicroTime 200 (PicoQuant, Berlin, Germany), a pulsed diode laser (LDH-P-C-470 PicoQuant, 470 nm, 20 MHz, 8 μW at the sample), a water immersion objective (1.2 NA, ×60, Olympus) mounted on a 3D piezo scanner (Physik Instrumente, Karlsruhe, Germany), and SPAD-MPD detector. The optics included Z473/635 RPC dichroic mirror and a 525/50 nm bandpass filter (Chroma Rockingham, VT, USA). SPBs labeled with fluorescent lipid analogue DOPE-Atto488 (1:100.000 dye:lipid) were visualized by scanning in their plane and the mean diffusion time of the dye was determined using Z-scan FCS, as previously described. [3] Briefly, fluorescence fluctuations caused by the lateral diffusion of individual dye molecules were recorded for 60 s at each point along the z-axis (with 200 nm steps) and the obtained data were fitted with appropriate parabolic functions gaining surface dye concentration and average lateral diffusion coefficient.

**Confocal Microscopy Imaging and Fluorescence Correlation Spectroscopy**

Imaging and FCS was performed on a home-built confocal microscope consisting of an inverted microscope bodyIX71 (Olympus, Hamburg, Germany). Pulsed picosecond laser (PicoTA532, PicoQuant, Berlin, Germany) was used at 25 MHz repetition rate. The output light of the optical fiber re-collimated by an air space objective (UPLSAPO 4X, Olympus, Hamburg, Germany). A quad-band dichroic mirror (ZT375/473/532/6–5rpc - Chroma Rockingham, VT, USA) was employed in order to up-reflect the light onto a water immersion objective (60x, NA 1.2, Olympus, Hamburg, Germany). Furthermore, a single photon avalanche diode (SPAD- MPD, Bolzano, Italy) using a 595/50 nm bandpass filter (Chroma Rockingham, VT, USA) was employed as a detection system.
For confocal microscopy, the power of the laser was kept below 2.5 μW, measured at the end of the fiber, and each image was recorded at a different resolution, depending on the size of the single GUV, scanning in a monodirectional mode. Before the measurements the ibidi chamber with glass bottom was coated with 200 μL of BSA-biot (0.1 g/L), 200 μL of streptavidin (2 mg/L), waiting 30 minutes for each step and afterwards washing each well with milliQ water. Each well was filled with buffer (10 mM Hepes, pH=7.4, 140 mM KCl, 0.1 mM EGTA, glucose; 300 mOsm $kg^{-1}$) and a GUV suspension was added and let to sediment for an hour. Subsequently, $CaCl_2$ stock solution was added to obtain a desired calcium concentration. CaM-R was added in the last step to get the final concentration of 30 nM. At least three images of three different GUVs were acquired for each well. Unless otherwise stated, measurements were performed after 4-8 hours of GUV incubation with CaM.
The same technical set-up was kept and an identical coating of the ibidi chambers with glass bottom was performed, except for the preliminary measurements of the labelled CaM in a buffer solution, in the absence of any GUV composition. In this case, the coating was achieved by using only Bovine Serum Albumine (BSA).

### Generalized Polarization

Steady-state emission spectrum ($\lambda_{ex}$=350 nm) and two excitation spectra ($\lambda_{em}$=440 nm and $\lambda_{em}$=490 nm) were recorded using FS5 spectrofluorometer (Edinburgh Instruments, UK). The excitation spectra were used to calculate excitation generalized polarization spectra that were averaged over excitation wavelengths to obtain mean *GP* values. *GP* was calculated using the following formula:

$$GP_{(\lambda_{ex})} = \frac{I_{440} - I_{490}}{I_{440} + I_{490}}$$

where $I_{440}$ and $I_{490}$ represent fluorescence intensities emitted at 440 nm and 490 nm, respectively (excited at the excitation wavelength $\lambda_{ex}$.[4] Titration experiments were performed in quartz cuvette ultra-micro cells purchased from Hellma Analytics (Plainview, NY USA) containing 20 µL of 60 µM CaM. The additions were done with increasing volumes in order to keep the CaM to lipid ratio between 1:1-1:100. To account for possible dilution effects, control experiment was performed in the same conditions but in the absence of CaM. Next, $CaCl_2$ was titrated to the cuvette containing 60 µM CaM and 50 µM LUVs. Direct effect of $Ca^{2+}$ was measured by performing control experiment in the absence of CaM. Since there is adsorption of $Ca^{2+}$ to the lipid bilayer, the GP in the presence of CaM ($GP_{CaM}$) was subtracted by the measured GP in the absence of CaM ($GP_0$), for each measured calcium concentration.The temperature was kept at 37±0.5 °C for all measurements.

### Simulation system construction and models: system, MD set-ups and analysis

All the tested systems were built using CHARMM-GUI platform, [5-6] using a total of 200 lipids [7-8] for each composition. The bilayers were solvated with water in a box of roughly 7.5 x 7.5 x12 nm$^3$ (x y z) size. A single molecule of Apo-CaM (calmodulin without any $Ca^{2+}$) was added to each lipid composition and placed in the bulk water. The systems were neutralized using KCl, and a further 0.15M of KCl was added to mimic cytosolic conditions. In the same way, we built 8 additional systems using Holo-CaM (calmodulin with 4 $Ca^{2+}$ in the binding sites). Yet another set of 8 additional systems was built taking the system containing Holo-CaM and adding $Ca^{2+}$ in the bulk water at the concentration of about 30 mM (see **Table S1** in the **Supplementary Material**). All these simulations were run in double replicates. Furthermore, for some of the systems we also built a set of systems containing Holo-CaM and 100-200mM of $Ca^{2+}$ ions (2 replicas each). Exact details about each system can be found in the **Supplementary Material**.

Simulations were performed using GROMACS 2020.x [9] package. The integration time step used to solve the equation of motion was 2 fs, and the Verlet lists algorithm for neighbor tracking [10]. Electrostatic interactions were treated using the smooth particle mesh Ewald (PME) [11-12]. Van der Waals interactions were treated using a Lennard-Jones potential with force-switching algorithm (1 nm–1.2 nm) and a cut-off of 1.2 nm. Simulations were run in the NpT ensemble and Nose-Hoover [13-14] thermostat and Parrinello-Rahaman[15] barostat were used to keep the temperature and the pressure constant at the values of 298 K and 1 atm, respectively. Bonds involving hydrogens were constrained with P-LINCS, [16-17] whereas the geometry of the used TIP3P model [18-19] was fixed using SETTLE [20].

The systems under analysis have a high charge density, e.g., Apo-CaM has a net charge of -24. Charge–charge intermolecular interactions are, therefore, likely more relevant than other non-

covalent interactions for the CaM–membrane interactions. Classical MD simulations and non-polarizable Force Fields are known for their limitations in describing systems with high charge density like in CaM containing systems [charge scaling manifesto: [21]]. For this reason, we applied Electronic Continuum Correction (ECC) to CHARMM36m Force Field,[22] which improves the description of charge-charge interactions [23-24].
All the analyses were performed discarding 1 μs (out of 5 μs). We calculated the binding percentage, defined as the fraction of the stored conformations where the protein is in contact with the membrane over time. The contacts were calculated using the gmx mindist tool from GROMACS, using a cutoff of 0.35 nm. Moreover, to shed light on the binding modes of the CaM on top of the membranes, we calculated the binding probability between each EF loop (named A, B, C, and D, see **Figure S1** in the **Supplementary Material**) and the membrane. After that, we calculated the probability of all the possible combinations of the binding of different EF loops at the same time, e.g., binding mode A means that only A is binding to the membrane, and binding mode ABCD means that all the 4 EF loops are binding at the same time to the membrane. The contact probability is the fraction (over time) where the protein is considered in contact with the membrane. The two systems are defined in contact if the distance between any protein atom and the membrane is below a properly chosen cut-off (0.35 nm). The label "apo"(first column from the left) refers to the bare protein, without any calcium ions added to the system. "Holo" (second column) refers to the protein with 4 calcium ions bonded to the calcium-binding sites and no free calcium in the solution. The last two columns (calcium 30 mM and calcium 150 mM) refers to the same system as "holo", but with additional calcium chloride in solution.

**Table S1.** Unbiased Molecular Dynamics Simulations Details. The total number of lipids is always 200. For binary lipid mixtures, e.g, A/B, the ratio between A and B is always 8:2. For ternary lipid mixtures, e.g., A/B/C, the ratio of each component is always 6:2:2.

| System | #Water[a] | #K+[b] | #Cl«[c] | #Ca[d] (bound) | #Ca[e] (solution) | #Replica |
|---|---|---|---|---|---|---|
| PC-ApoCaM | 16676 | 68 | 44 | 0 | 0 | 2 |
| PE-ApoCaM | 14410 | 61 | 37 | 0 | 0 | 2 |
| PE/PC-ApoCaM | 14796 | 63 | 39 | 0 | 0 | 2 |
| PC/PS-ApoCaM | 16259 | 107 | 43 | 0 | 0 | 2 |
| PE/PS-ApoCaM | 14510 | 102 | 38 | 0 | 0 | 2 |
| PC/CH-ApoCaM | 15380 | 65 | 41 | 0 | 0 | 2 |
| PE/CH-ApoCaM | 13617 | 59 | 35 | 0 | 0 | 2 |
| PE/PC/CH-ApoCam | 14017 | 61 | 37 | 0 | 0 | 2 |
| PC/PS/CH-ApoCaM | 14937 | 104 | 40 | 0 | 0 | 2 |
| PE/PS/CH-ApoCaM | 13637 | 99 | 35 | 0 | 0 | 2 |
| PC-HoloCaM | 16764 | 60 | 44 | 4 | 0 | 2 |
| PE-HoloCaM | 14418 | 53 | 37 | 4 | 0 | 2 |
| PE/PC-HoloCaM | 14803 | 55 | 39 | 4 | 0 | 2 |
| PC/PS-HoloCaM | 16276 | 99 | 43 | 4 | 0 | 2 |
| PE/PS-HoloCaM | 14518 | 94 | 38 | 4 | 0 | 2 |
| PC/CH-HoloCaM | 15388 | 57 | 42 | 4 | 0 | 2 |
| PE/CH-HoloCaM | 13625 | 51 | 35 | 4 | 0 | 2 |
| PE/PC/CH-HoloCaM | 14025 | 53 | 37 | 4 | 0 | 2 |
| PC/PS/CH-HoloCaM | 14945 | 96 | 40 | 4 | 0 | 2 |
| PE/PS/CH-HoloCaM | 13645 | 91 | 35 | 4 | 0 | 2 |
| PC-HoloCaM | 16764 | 40 | 44 | 4 | 10 | 2 |
| PE-HoloCaM | 14418 | 33 | 37 | 4 | 10 | 2 |
| PE/PC-HoloCaM | 14803 | 35 | 39 | 4 | 10 | 2 |
| PC/PS-HoloCaM | 16276 | 79 | 43 | 4 | 10 | 2 |
| PE/PS-HoloCaM | 14518 | 74 | 38 | 4 | 10 | 2 |
| PC/CH-HoloCaM | 15388 | 37 | 41 | 4 | 10 | 2 |
| PE/CH-HoloCaM | 13625 | 31 | 35 | 4 | 10 | 2 |
| PE/PC/CH-HoloCaM | 14025 | 33 | 37 | 4 | 10 | 2 |

| | | | | | | |
|---|---|---|---|---|---|---|
| PC/PS/CH-HoloCaM | 14945 | 76 | 40 | 4 | 10 | 2 |
| PE/PS/CH-HoloCaM | 13645 | 71 | 35 | 4 | 10 | 2 |
| PC/PS-HoloCaM | 16147 | 79 | 123 | 4 | 50 | 2 |
| PE/PS-HoloCaM | 14398 | 74 | 118 | 4 | 50 | 2 |
| PC/PS/CH-HoloCaM | 14825 | 76 | 120 | 4 | 50 | 2 |
| PE/PS/CH-HoloCaM | 13525 | 71 | 115 | 4 | 50 | 2 |

[a] Number of water molecules [b] number of potassium ions [c] number of chloride ions [d] the number of calcium covalently bound to calmodulin EFs loops [e] number of calcium added to the bulk water.

### Chemical structures of lipids used in the experiments

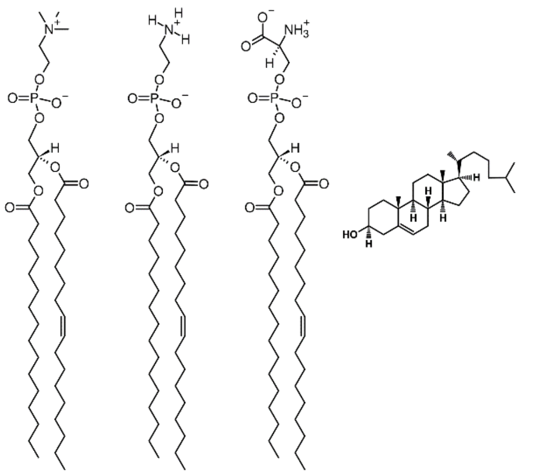

**Figure S1.** Chemical structures of the components of model lipid membranes used in this study. From left to right: POPC (1-palmitoyl-2-oleoyl-sn-glycero-3-phosphocholine), POPE (1-palmitoyl-2-oleoyl-sn-glycero-3-phosphoethanolamine), POPS (1-palmitoyl-2-oleoyl-sn-glycero-3-phospho-L-serine), and cholesterol.

### Amino acids and DNA sequences of the various calmodulin used in the experiments

Wheat calmodulin primary sequence employed for fluorescence techniques
MADQLTDDQIAEFKEAFSLFDKDGDGCITTKELGTVMRSLGQNPTEAELQDMINEVDADGNGTIDFPEF
LNLMARKMKDTDSEEELKEAFRVFDKDQNGFISAAELRHVMTNLGEKLTDEEVDEMVREADVDGDGQ
INYDEFVKVMMAK

Bovine calmodulin primary sequence used for label-free techniques
MADQLTEEQIAEFKEAFSLFDKDGDGTITTKELGTVMRSLGQNPTEAELQDMINEVDADGNGTIDFPEF
LTMMARKMKDTDSEEEIREAFRVFDKDGNGYISAAELRHVMTNLGEKLTDEEVDEMIREADIDGDGQV
NYEEFVQMMTAK

Porcine calmodulin primary sequence used for label-free techniques
MADQLTEEQIAEFKEAFSLFDKDGDGTITTKELGTVMRSLGQNPTEAELQDMINEVDADGNGTIDFPEF
LTMMARKMKDTDSEEEIREAFRVFDKDGNGYISAAELRHVMTNLGEKLTDEEVDEMIREADIDGDGQV
NYEEFVQMMTAK

DNA Sequence of calmodulin employed for live cell experiments
ATGGCTGACCAGCTGACTGAGGAGCAGATTGCAGAGTTCAAGGAGGCCTTCTCCCTCTTTGACAAGGA
TGGAGATGGCACTATCACCACCAAGGAGTTGGGGACAGTGATGAGATCCCTGGGACAGAACCCCACTG
AAGCAGAGCTGCAGGATATGATCAATGAGGTGGATGCAGATGGGAACGGGACCATTGACTTCCCGGAG

TTCCTGACCATGATGGCCAGAAAGATGAAGGACACAGACAGTGAGGAGGAGATCCGAGAGGCGTTCCG
TGTCTTTGACAAGGATGGGAATGGCTACATCAGCGCCGCAGAGCTGCGTCACGTAATGACGAACCTGG
GGGAGAAGCTGACCGATGAGGAGGTGGATGAGATGATCAGGGAGGCTGACATCGATGGAGATGGCCAG
GTCAATTATGAAGAGTTTGTACAGATGATGACTGCAAAGTGA

### Results and Discussion

#### Live cell imaging

We have used CaM fluorescently-tagged with EGFP (EGFP-CaM) and focused on membrane-rich surface protrusions visible at the edge of transiently transfected HeLa cells. In a resting state, little to no signal of EGFP-CaM was detected in membrane protrusions (**Figure S2A**, 0 min, first row). On the contrary, a strong and fairly homogeneous EGFP-CaM signal was found in the cell body. Treatment of cells with ionomycin, which induces release of $Ca^{2+}$ from intracellular stores and increases cytosolic $Ca^{2+}$ concentration, increased the presence of EGFP-CaM therein (**Figure S2A**, 8 min, first row). **Figure S2A** shows the images acquired each 2 minutes after addition of ionomycin (0 and 8 minutes images were already shown in the main text). No measurable changes were observed in the cell body. Quantification of relative EGFP-CaM intensities (**Figure S2B**) in membrane protrusions indicates its significant accumulation therein after treatment of cells with 0.2 μM ionomycin. These data indicate that increased intracellular $Ca^{2+}$ levels cause partial CaM relocalization into membrane-rich environment of surface protrusions. Our results indicate that under certain conditions, such as calcium intake, CaM and membranes can be functionally connected.

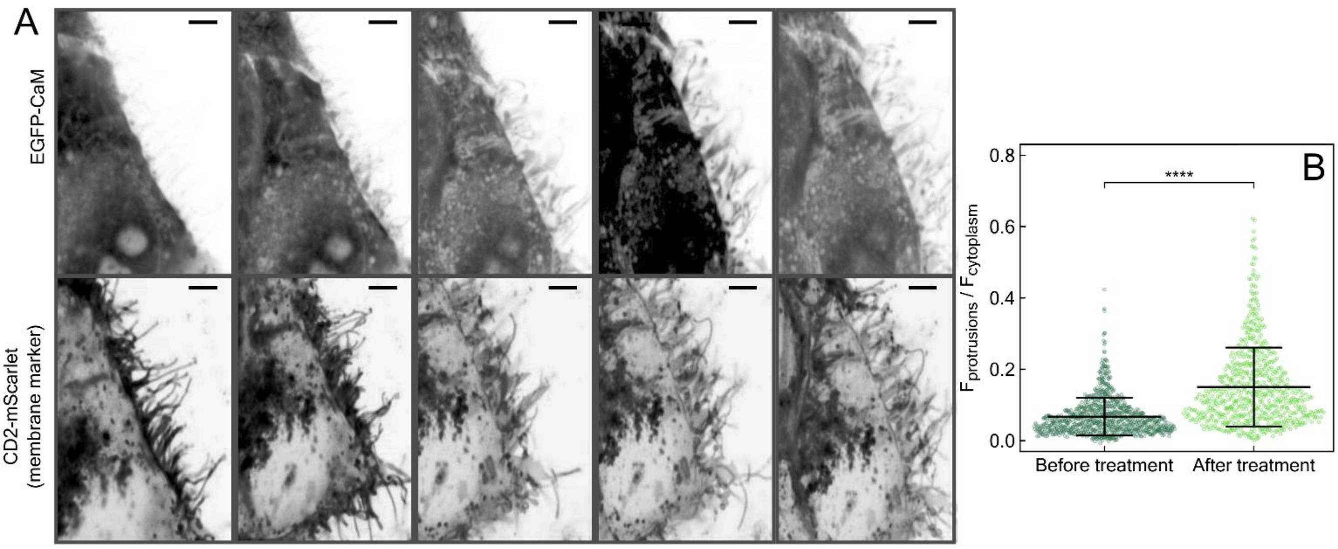

**Figure S2**. a) Time-lapse live-cell imaging of transiently transfected HeLa cells expressing EGFP-CaM in the cytosol and CD2-mScarlet at the plasma membrane demonstrating EGFP-CaM relocalization to membrane protrusions after the treatment of cells with 1 μM ionomycin (upper). Changes in morphology of membrane protrusions were visualized using a membrane marker (CD2-mScarlet; lower). Scale bars of 5 μm are shown in the right upper corners. b) The ratio of EGFP-CaM fluorescence intensity in the protrusions and in the cytoplasm before and after the treatment of cells with 0.2 μM ionomycin (Before treatment: N(cells) = 118; N(protrusions) = 886; After treatment: N(cells) = 98; N(protrusions) = 686). Values represent normalized signal measured in protrusions correlated to the cytosolic EGFP-CaM levels (see **Experimental Procedure** for details). Significance of the results was tested using t-test (****, $p < 0.0001$).

### Characterization of the SPB via z-scan FCS

It is noteworthy that the presence of 60 mol% of PE considerably reduced stability of SPB membranes and prevented a more systematic SPR measurements. Only about 70% of SPR measurements with lipid composition containing 60 mol% PE could be analyzed. The instability of the studied SPBs was studied using fluorescence confocal microscopy imaging and Z-scan fluorescence correlation spectroscopy (FCS) performed on the support used for SPR measurements (see **Experimental Procedure**). These fluorescence experiments showed a success rate for PE/PS/CH SPBs formation of about 70%. Moreover, the obtained SPBs were inhomogeneous, i.e., contained adsorbed vesicles and defects, which could influence their interactions with CaM.

### Fluorescence Correlations Spectroscopy (FCS)

We performed single-point FCS measurements directly after 30 minutes from the addition of CaM-R to the chamber. As schematically illustrated in Figure S3 we measured CaM-R diffusion at the GUV membrane and compared it with its diffusion in bulk solution.

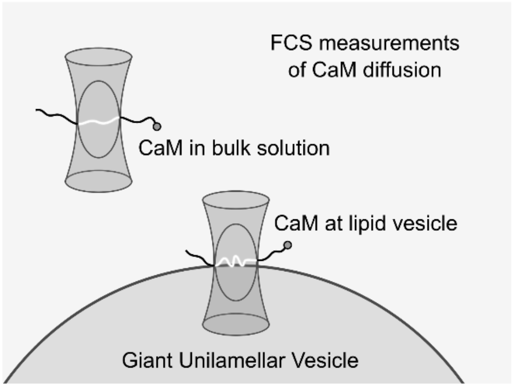

**Figure S3**. Schematic cartoon of the FCS measurements of calmodulin labeled with rhodamine B (CaM-R) in solution and at the top of the GUV membrane.

To measure the diffusion time of CaM-R in the bulk, one or a few single points in solution were measured for 2-5 minutes. The autocorrelation curves were fitted using SymPhoTime 64 software, applying a gobal analysis for multiple measurements at once, selecting a 3D free diffusion model, single species:

$$G_{(t)} = \frac{1}{N}\frac{1}{1+\left(\frac{t}{\tau_b}\right)}\sqrt{\left(\frac{1}{1+\left(\frac{t}{\tau_b}\right)\kappa^2}\right)}$$

where $N$ is the number of independently diffusing species within the confocal volume, $\tau_b$ is the mean diffusion time in the bulk, and $\kappa$ is the structural parameter describing the shape of the confocal volume. [25-28] Size of the confocal volume and $\kappa$ were calculated for aqueous solution of rhodamine-B as a standard ($D$ = 427 ± 4 µm$^2$/s at 298.15 K). [29] The mean diffusion time of the free bulk CaM-R determined this way was $\tau_b$ = 0.20 ± 0.04 ms. Diffusion coefficients ($D$) were calculated from diffusion times ($\tau$) according to:

$$\boldsymbol{D} = \frac{\boldsymbol{\omega_0}^2}{4\boldsymbol{\tau}}$$

where $\omega_0$ is the waste of the confocal volume calculated using rhodamine-B, as described above. The volume of the confocal volume was 0.77 ± 0.02 fL. For our $\tau_b$, $D$ = 100 ± 20 µm$^2$·s$^{-1}$, which is a typical value for the diffusion of a small protein in water at 25 °C.

For CaM-R diffusion at the lipid membrane, at least 10 different GUVs in each chamber were measured by recording the intensity fluctuations at the their surface for 1 minute. We performed the measurements 30-90 min after CaM addition into the chamber. We performed a global fitting for all the measurements performed for the same lipid composition (separately for the samples measured in the absence and in the presence of 10 mM $CaCl_2$) using SymPhoTime 64 software. We used two-component model:

$$\boldsymbol{G}_{(t)} = \left[\frac{\boldsymbol{\rho_b}}{1+\left(\frac{t}{\tau_b}\right)}\sqrt{\left(\frac{1}{1+\left(\frac{t}{\tau_b}\right)\kappa^2}\right)} + \frac{\boldsymbol{\rho_m}}{1+\left(\frac{t}{\tau_m}\right)}\sqrt{\left(\frac{1}{1+\left(\frac{t}{\tau_m}\right)\kappa^2}\right)} + \right]$$

where $\tau_b$, $\tau_m$, $\rho_b$ and $\rho_m$ are the diffusion times and amplitudes of the two different diffusing species, i.e., CaM in solution and CaM at the GUV membrane. The sum of the amplitudes is proportional to the inverse of $N$. [25-28] We fixed $\tau_b$ at 0.2 ms, as determined above, and treated $\tau_m$ and $\rho_m$ as unrestrained parameters; for $\rho_m$ 0.002-1000 ms boundaries were set.

When focused on the top surface of a GUV, a second significantly slower diffusion fraction occurred. The results are summarized in Figure 2A (main text), where we show the longer diffusion time, its fraction, and the associated standard error for PC, PC/PS, PE/PS/CH, and PE/PC/PS/CH, in the absence and in the presence of 10 mM $CaCl_2$. For PC and PC/PS a fraction of <30% with diffusion time of 5-10 ms ($D$ = 4-2 µm$^2$·s$^{-1}$) was found. Adding PE to the lipid mixture (i.e., for PE/PS/CH and PE/PC/PS/CH) increased that faction to 40-70% with diffusion times of about 8-10 ms ($D$ = 3-2 µm$^2$·s$^{-1}$). Although these values are in the range typical for 2D protein diffusion,[30] their large uncertainty hampers further interpretation. Addition of 10 mM $CaCl_2$ further increased the contribution of the slow diffusing components for these PE-containing GUVs, reaching a diffusion time of ~25 ms for PE/PS/CH. FCS shows that even in the absence of PE and $Ca^{2+}$ there is a small fraction of slower diffusin CaM. However, due to the presence of POPE and calcium the fraction of the slow diffusion component increases more than 3 times confirming that these components are essential for the CaM-membrane interactions.

### Supplementary MD data

Figure S4 shows how CaM binds to membrane of different compositions and at different $Ca^{2+}$ concentrations. EF loops are named as A (amino acids (aas) 10–40), B (aas 45–75), C (aas 82–112) and D (aas 118–148), and visually represented in Figure S4 by blue, red, yellow, and green colours, respectively. The contact probability was calculated for the "apo", "holo" at 30 mM and 150 mM calcium (see Table S1 for simulation details). Different binding modes are classified based on which EF loops (one or more) bind to the membrane surface, e.g., A means that only EF loop "A" is binding, whereas "A, B, C, D" means all 4 EF loops are binding at the same time. Overall the most frequent binding modes are the ones where only one EF loop at the time binds the membrane, regardless the composition. Only for systems containing PE and $Ca^{2+}$ in solutions, binding modes with more EF loops bound at the same time are populated, e.g,, the binding mode "AB" and "ABCD". This tendency disappears when PS is added to membranes containing PE, indicating that the presence of PE increases the chances for the protein to lie down on the membrane surface, compared to other lipid compositions.

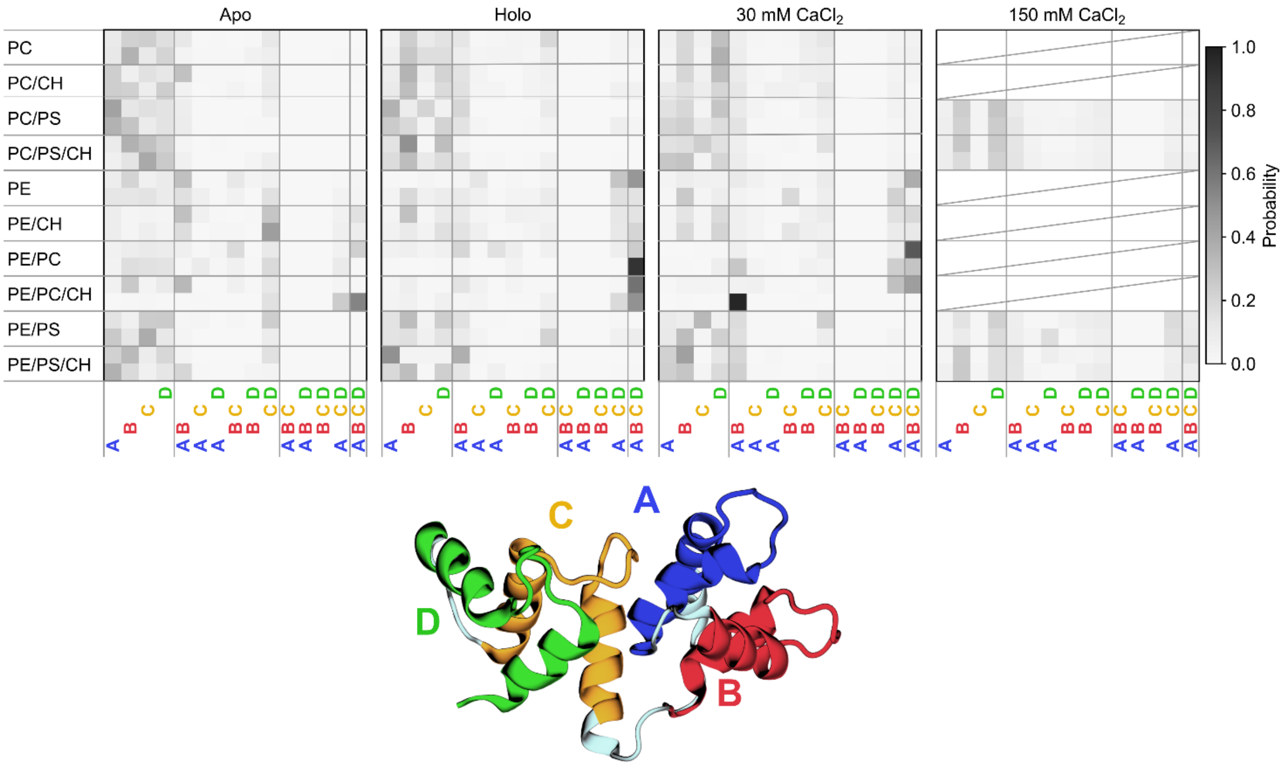

**Figure S4.** Contact probability maps between EF loops of CaM and lipid membranes with different compositions. From left to right the contact maps are reported for the apo-CaM, holo-Cam, holo-CaM at 30 mM $CaCl_2$, and holo-CaM at 150 mM $CaCl_2$. The X-axis labels of the contact maps represent all the possible combinations for the EF loops binding at the same time. EF loops are named as A, B, C and D, as shown in the CaM structure depicted in the figure. For each system two replicates were simulated and the probabilities for them are shown in two separate rows for each lipid composition.
